# Direct empirical in-house assessment of peptide proteotypicity for targeted proteomics

**DOI:** 10.64898/2026.02.22.699713

**Authors:** IO Butenko, NA Kitsilovskaya, IK Chudinov, AV Vakaryuk, AA Lebedeva, NM Baraboshkin, AG Khchoian, AV Kovalenko, VD Gremyacheva, AA Lazareva, OV Kurylova, KS Gorbunov, AV Pavlenko, GL Kozhemyakin, OV Fedorov, EN Ilina, VM Govorun

## Abstract

In bottom-up proteomics peptide it was early shown that despite a certain protein is present in a sample, only a subset of it’s proteolytic peptide products will be detected with LC-MS analysis. Property of peptide being frequently detected given its source protein’s identification was called proteotypicity. Much effort has been since applied to predict proteotypic peptides and summarize evidence on peptide detection. Nevertheless, when targeted proteomics method is being developed, prediction or inference from communal experience might be inaccurate and prior knowledge of true peptide proteotypicity in a selected setup for a selected population is necessary. In this work we test fully in-house approach for proteotypicity assessment including comprehensive peptide synthesis and detection verification. Proteotypicity and contribution of sample processing and biology-related factors are estimated in a model experiment for three plasma proteins, albumin, ceruloplasmin and C-reactive protein.

## Introduction

Targeted bottom-up proteomics is a method of choice for second step in a triangular strategy biomarker discovery [1]. It is typically performed with a relatively high-throughput UHPLC separation and triple-quadrupole detector operating in a MRM mode. MRM acquisition performed with triple quadrupole instruments is the most sensitive detection method when instruments of the same generation are compared with the same ionization technique. In addition, it is much less prone to data omission when compared to DDA or DIA acquisition and provides comparable specificity of detection [2, 3]. MRM targeted at pairs of naturally abundant tryptic peptides produced during tryptic digestion and their analogues synthesized with C-terminal lysine or arginine labelled with stable isotopes used as internal standards allows to achieve precise and accurate quantitation of peptides. Another approach relies on recombinant isotopically labelled proteins (QConCat and QPrest technologies) that are added to the samples at the early stages and produce internal standard peptides during tryptic digestion.

All targeted methods require prior knowledge of peptides to target, time and effort are required to build a quantitative method for each target and in case of misselection of a target peptide, used for quantitation of a protein of interest, time and cost of producing standards are lost at least partially for a multiplex protein standard and completely for a synthetic peptide. Selection of an unsuitable peptide can lead to an inconsistent detection of protein in a sample where a protein is present, thus leading to false negative results and compromising biomarker discovery or validation. Checking the detection of a peptide in a reference sample or with a pure protein sample might be not enough to ensure consistent detection. This is a long-known phenomenon, observed at the early days of bottom-up proteomics.

Reproducible detection of a certain protein of interest has been a cornerstone question in the development of bottom-up proteomics. It was early observed that only a fraction of theoretically predicted peptides that can be produced by proteolysis of proteins by trypsin is in fact detected by mass-spectrometry (i.e. sequence coverage for any identified protein is almost never complete) experiment and this observation still holds after 20 years of technology development [1], [2]. To address the phenomenon of experimental tryptic peptide observation or a lack of one, two groups have proposed a term “proteotypic peptide” for reproducibly detectable peptides in 2005. Both groups have used this term for experimentally observable peptides in a bottom-up shotgun proteomics experiment. Kuster and Aebersold et al. have defined a proteotypic peptide as “an experimentally observable peptide that uniquely identifies a specific protein or protein isoform”, uniqueness requirement is necessary for the task of quantitative proteome scoring proposed in this seminal paper. Beavis, Cortens and Craig define proteotypic peptide as “those peptides in a protein sequence that are most likely to be confidently observed by current MS-based proteomics methods”. While peptide uniqueness (or, more strictly peptide sequence specificity for a product of certain protein-coding gene) is extremely important for straightforward interpretation of proteomic profiling results in most applications, the definition given by Beavis, Cortens and Craig is more focused on a problem of complex influence of multiple methodological factors on peptide detection. Their definition is closely related to a question of whether any theoretically observable peptide is in fact fit for the purpose of protein detection, quantification and further comprehensive quantitative profiling of sample collections by a certain selected method as required for example in circulating plasma protein biomarker studies.

Many attempts were made to predict proteotypic peptides or estimated probability of peptide identification for a protein or proteome of interest based on either protein sequences or deposited bottom-up proteomics data. Several approaches have immediately emerged for prediction of proteotypic (also “signature”, “best flyer”, “flyable”, “observable” or “high-responding”, “enhanced signature”, if suitable for protein quantitation - “quantotypic”) peptides. PeptideAtlas project estimates proteotypicity of a peptide in a given sample type (e.g. human plasma) based on identifications from hundreds of experiments performed on a wide variety of technological platform and multiple species and tissue specific peptide databases have been created with a narrower scope. Multiple attempts in proteotypicity prediction showed that predictive models work well for a technological platform (complete sample preparation technique, separation and detection methods) and for the same sample type (biological fluid, tissue or cell culture).

Thus, development of quantitative targeted bottom-up proteomics method for the purpose of biomarker discovery and analysis of large sample cohorts can possibly significantly benefit from prior direct estimation of selected peptide proteotypicity for a specific technological platform and population. In this work we apply comprehensive peptide synthesis for three major circulating plasma proteins and assess peptide proteotypicity based on a set of plasma samples obtained from 195 patients, that belong to the general population that is expected to be tested, and processed with a define sample preparation and detection procedures to be used as a technological solution for quantitation of more proteins of interest. We compare are results with PeptideAtlas estimation for proteotypicity and several predictions made by other groups. We observe that relying solely on prediction or aggregated data leads to both selection of peptide with low proteotypicity and omission of higly proteotypic peptides.

## Materials and Methods

### Peptide selection

Human protein info for downloaded from neXtProt database (<u>10.1093/nar</u>/g<u>kz995</u>, <u>10.5281/zenodo.14163587</u>) and processed with in-house R script using magrittr, httr2, dplyr, readr, tidyr, stringi, Peptides, pbapply, parallel, data.table and ggplot2 packages. Code is avalable in Supplementary Material 1. Tryptic peptide sequences for every protein were obtained and peptides were scored based on their gene-wise uniqueness and proteoform coverage, amino acid composition, physico-chemical properties and position in the protein chain.

### Peptide synthesis

Peptide were synthesized with solid-phase chemistry in batches according to their length (see Supplementary Table).

#### Chorus

Peptides were synthesized according to the solid-phase peptide synthesis protocol on the PurePep Chorus automated peptide synthesizer (Protein Technologies, Inc.) using 9-fluorenylmethylmethoxycarbonyl (Fmoc)-protected amino acid derivatives. N-alpha-Fmoc-Ng-(2,2,4,6,7-pentamethyldihydrobezofuran-5-sulfonyl)-L-arginine-Wang resin and N-alpha-Fmoc-N-epsilon-Boc-L-lysine-Wang resin were used as the solid phases. N,N’-Diisopropylcarbodiimide (DIC) in the presence of Ethyl cyano(hydroxyimino)acetate (Oxyma Pure) was used for Fmoc-amino acid condensation. A 15-fold excess of Fmoc-amino acids relative to the carrier capacity was used in the addition reaction. Removal of the protective Fmoc groups from the first amino acid on the carrier and from the growing peptide chain was performed with a 20% solution of 4-methylpiperidine (v/v) in N,N-dimethylformamide. The N-terminus was capped using a 50-fold excess of 5% propionic anhydride (v/v) in DMF.

#### Liquid handler-based plate synthesis

Peptides were synthesized according to the solid-phase peptide synthesis protocol on the automated liquid handler system Freedom EVO (Tecan) using 9-fluorenylmethylmethoxycarbonyl (Fmoc)-protected amino acid derivatives. N-alpha-Fmoc-Ng-(2,2,4,6,7-pentamethyldihydrobezofuran-5-sulfonyl)-L-arginine-Wang resin and N-alpha-Fmoc-N-epsilon-Boc-L-lysine-Wang resin were used as the solid phases 2-(1H-Benzotriazole-1-yl)-1,1,3,3-tetramethylaminium tetrafluoroborate (TBTU) in the presence of 1-Hydroxybenzotriazole (HOBt) and N,N-Diisopropylethylamine (DIPEA) was used for Fmoc-amino acid condensation. A 15-fold excess of Fmoc-amino acids relative to the carrier capacity was used in the addition reaction. Removal of the protective Fmoc groups from the first amino acid on the carrier and from the growing peptide chain was performed with 5% piperazine + 1% DBU + 1% FA in N,N-dimethylformamide.

#### Cleavage from resin and unblocking of side groups

Removal of peptides from the solid phase and simultaneous unblocking of side groups was carried out in a mixture of trifluoroacetic acid : 3,6-dioxa-1,8-octandithiol : triisopropylsilane : anisole : water in a volume ratio of 183:5:2:5:5 for 3 hours. At the end of incubation, the mixture was withdrawn and cooled diethyl ether was added to the resulting solution. The resulting suspension was cooled for 30 min at −20°C and then centrifuged for 10 min, at 7000 rpm. The resulting precipitate was lyophilic dried and then dissolved in 1 mL of 5% acetonitrile solution in water for purification.

### Collection description

Blood samples were collected in tubes containing anticoagulant (ethylenediaminetetraacetic acid) and immediately centrifuged at 3500 rpm for 5 min to separate plasma. The separated plasma was stored at −20 °C until measurement. The study was conducted in accordance with the Declaration of Helsinki.

#### Healthy volunteers

In total, 96 samples of healthy individuals (66 females and 30 males, 42-75 years, median - 62 years) plasma were used for proteotypicity estimation. This study involving humans was approved by the institutional review board of FSBEI of Higher Education Northern State Medical University of the Ministry of Health of the Russian Federation (approval number: 07/09-11 from 28/09/2022).

#### ICU patients

Blood samples of 10 patients (6 males, 4 females, 46-76 years, median - 60 years) were collected daily (4-19 days per patient, median - 7 days) during their stay in the ICU. C-reactive protein was measured within clinical laboratory using immunoturbidimetry on a Beckman Coulter biochemical analyzer for 91 of 96 samples (4.9 - 256.3 mg/L, median - 90 mg/L). This study involving humans was approved by the institutional review board of FSAI NMRC “Treatment and Rehabilitation Center” of the Ministry of Health of the Russian Federation (approval number: EC/052 from 04/12/2023). The study was conducted in accordance with the Declaration of Helsinki.

### Sample preparation

Batches of 80-96 plasma samples were processed in a 96-well plate with Tecan Freedom EVO 150/8 liquid handling station in a single batch. Plasma samples were assigned random positions in a plate, 10 µl of each sample was transferred into a well plate and processed. In the first step samples were peptized with urea (6 M) and disulfide bonds were reduced with dithiothreitol (13 mM) during 30-minute incubation at 37°C, in the second step free thiol groups of cysteines were alkylated with freshly prepared iodoacetamide (40 mM) during 30-minute incubation at room temperature in the dark. Both reactions were performed in a presence of 300 mM tris adjusted to pH 8.0 with hydrochloric acid. After that samples were diluted with the same buffer solution to drop urea concentration down to 0.55 M. 14 mg of bovine trypsin in 1 M calcium chloride prepared immediately before addition by in-house procedure to be published later was added to samples so that estimated weight ratio of protein to protease was 50 to 1 and calcium chloride concentration was 3.5 mM. After 1-hour incubation at 37°C the same amount of bovine β-trypsin (produced in-house from crystallized trypsin, Samson Med, Russia) solution was added, so that estimated protein to protease became 25 to 1 and calcium chloride concentration - 7 mM. Samples were incubated overnight (at least 18 hours). After incubation samples were diluted with trifluoracetic acid (TFA) and acetonitrile (ACN) solution so that final urea concentration was 150 mM, TFA - 1% volumetric and acetonitrile – 5%. Final sample volume was 1400 µl.

Peptides were extracted by solid-phase extraction using 96-well plate with C18 for desalting and Tecan Resolvex A200 positive pressure processor. All solvents used as a mobile phase were modified with 0.2% TFA and volumes of 600 µl were used for each step. The plate was conditioned with three volumes acetonitrile and equilibrated with three volumes of 5% ACN. Sample digests were loaded and wells were washed three times with 5% ACN, and the bound peptides were eluted with a volume of each 30%, 50% and 80% acetonitrile. Eluates were combined and dried using a speed vacuum concentrator. Plasma samples and calibrants were then resolubilized in 35 µl of 5% ACN with 0.1% TFA.

### LC-MS

#### Separation conditions

Сhromatographic separation of peptides was performed by reversed-phase chromatography using a Ultra AQ C18 column (C18, 2.1 × 150 mm, 3 μm, Restek, USA) at 30°C with a flowrate of 300 μl/min. Mobile phase solvents A (5% ACN) and B (80% ACN) were modified with 0.1% formic acid. Gradient went from 5% B to 18.5% in 0.5 min, to 32% in 4.5 min, to 47% in 3 min, to 100% for 0.5 min and back to 5% for 2/5 min.

Spray voltage was 5500 V, source temperature – 350°C, drying and nebulizer gas flow were set to 50 L/min and 30/L min. Mass-spectrometer acquired scheduled MRM data in the positive mode. Acquisition cycle was set to 400 ms with 90 s detection windows for each peptide.

#### Targeted method development

Crude peptide samples were transferred to 96-well plate and analyzed with 5 uL injections. Skyline (<u>10.1002</u>/<u>pmic.201200042</u>) was used to produce all theoretical product ions of b and y series with charge +1 for parent ions charged +2 and +3. For cysteine-containing peptides a set of transition was produced also for fully carbamidomethylated peptide and tested on a DTT and IAA-treated crude samples besides raw one. Treatment conditions were the same as for plasma samples.

First run was unscheduled and included declustering potential (DP) variation ( <u>10.1021</u>/<u>pr801122b</u>) while using predicted collision energy (<u>10.1021</u>/<u>pr1004289</u>). First run was used to determine retention time and optimal DP. Second run was performed in a scheduled manner with optimal DP and with the same approach to collision energy variation. Second run was used to select optimal CE, charge state and a set of 3-5 most intense fragment ions. Both DP and CE were optimized per precursor ion (not per individual fragment). Determined RT, DP, CE, charge state and transition set were used for each peptide in further analysis.

#### Plasma sample analysis

Samples were analyzed with 10 µl injection, thus equivalent of 2.85 ul of plasma was injected. MRM was performed in a scheduled manner. Acquisition cycle was set to 400 ms with 90 s detection windows for each peptide. Transition list included 100-613 transitions depending on a set of peptides tested. Plasma samples were analyzed in sets of 80-96 samples. Each testing batch included single plasma set and a peptide set. Each batch was accompanied with iRT injection and synthetic peptide pooled sample before the batch, at least once during the batch analysis and after the batch, followed by 5 blank injections that were manually inspected not to produce peptide detection.

### Data processing

Raw data was imported into Skyline, where automated peak detection was performed followed by manual inspection and for pooled synthetic standard sample runs, blank runs and at least 12 plasma runs. Table, containing results for individual fragment was processed using in-house R script (available in Supplementary Material).

### Data availability

## Results and Discussion

### Synthesis

Two strategies for peptide synthesis were tested. The first one utilized commercially available PurePep Chorus four-channel synthesizer and the second one utilized Tecan Freedom EVO liquid handler that was used to synthesize peptides in batches of 48. Both approaches allowed to produce 48 crude peptide samples per week. Chorus-based approach required 1 trained chemist and 2 lab assistants working in accurately scheduled shifts to synthesis twice a day with at least 4 hours hands-on time for each person daily. Peptides had to be split in pairs of batches, one batch for short synthesis during the day, and one batch for long overnight synthesis with total lab day duration was about 12 hours. Liquid-handler based approach required single launch a week, 1 trained chemist and 1 lab assistant working with typical schedule and about 4 hour hands-on time three times a week for the same output of 48 crude peptide samples. It is also easily scaled to three 48-peptide syntheses a week, which requires, though, another lab assistant to perform peptide cleavage after synthesis. In addition, liquid handler based approach was successful at lower scale, so cost per peptide detectability test was lower, which is important during the earlier stage of peptide screening.

### Method development for comprehensive peptide screening

Three plasma proteins, albumin (P02768), ceruloplasmin (P00450) and C-reactive protein (P02741) were selected for comprehensive peptide synthesis. All three proteins circulating in blood plasma are primarily produced by liver hepatocytes and excreted into bloodstream. In healthy individuals concentrations of these proteins are reported to be about 0.2 pmol per uL of plasma for C-reactive protein, 1.3 pmol/uL for ceruloplasmin and 0.5 nmol/uL for albumin (3.4 mg/L, 0.16 g/L and 40 g/L respectively). While albumin and ceruloplasmin exhibit relatively narrow range of concentrations in human blood plasma, C-reactive protein is a known acute phase protein and detected range spans at least two orders of magnitude. These proteins were selected to study peptide proteotypic properties unbiased by limit of detection.

Albumin has 2 shortened spliceforns described and detected at protein level and C-reactive protein has one and ceruloplasmin has no isoforms produced by alternative splicing. For each protein *in silico* proteolysis by trypsin was performed for all isoforms expecting complete and specific hydrolysis of peptide bond only C-terminal to lysine and arginine residues but not N-terminal to proline residues ( ). All peptides were selected for synthesis excluding C-terminal peptides, peptides that are not gene-specific (i.e. not “unique” to the protein) when all proteoforms of all other human genes are considered and peptides exceeding 30 residues in length.

C-terminal peptides were excluded for these proteins due to synthesis procedure limitations as they do not bear C-terminal lysine or arginine residues. This requires acquisition or production of separate solid phase for synthesis, including isotopically labelled residues for further analysis. This increases the cost of production of such peptides. While peptides that lack C-terminal lysine or arginine are typically rarely detected due to lower ionization efficiencies and protein termini are prone to degradation, they are typically avoided in targeted quantitative proteomics efforts.

Non-gene-specific peptides were mostly relatively short (55 peptides of 4 residues or less and 6 peptides of 5 or 6 residues). Shorter peptides area typically relatively hydrophilic and special care has to be taken to ensure separation conditions that allow reproducible and quantitative detection as they are often unretained during solid-phase extraction or LC-MS analysis. In addition, low mass range is prone to higher baseline in complex biological samples, thus short peptides are typically avoided in targeted proteomics. With the length of peptides duration of synthesis increases and synthesis efficiency decrease and as these peptides are rarely detected in bottom-up proteomics, we excluded peptides above 30 residues in length.

In total, synthesis of 103 peptides ranging in length from 5 to 30 residues was attempted. Peptide information is available in Supplementary Material 2. 30 peptides were proteoform-specific, 33 contained cysteine residues, 14 - tryptophan and 57 - neither of two. Three shorter peptides (5-6 residues, hydrophobicity −1.7, −1 and −0.4 on Kyte-Doolittle scale), three longer peptides (22, 23 and 30 residues, all having two asparagine residues, repeats and tryptophan residues) and 5 peptides bearing either several cysteine residues or tryptophan residue or 2 or 3-residue repeats did not allow a detection method to be established due to lack of expected peak group in an LC-MS MRM analysis with all predicted ions as described in Method section. All these sequence features are know to induce problems during SPE peptide synthesis and LC-MS analysis. In total, detection methods were developed for 93 peptides (10 for C-reactive protein, 56 and 57 for ceruloplasmin and albumin), including additional detection methods for 30 cystein-bearing peptides in a fully carbamidomethylated form, in total MRM method for multiplex LC-MS detection of 123 moieties was developed.

### Comprehensive peptide screening on individual plasma samples

The developed method was tested on two batches of individual (not pooled) plasma samples. First, peptide detection accuracy was assessed, than peptide detection rate was determined for every peptide and compared to Empirical Proteotypic Score reported in latest PeptideAtlas Human Plasma build (2025-08) and peptide properties.

#### Peptide detection assessment

To ensure correct peptide detection with Skyline peak detection algorithm, several runs of pooled peptide standards were LC-MS results were inspected manually, 36 runs with 123 MRM ion chromatogram groups each (4428 groups totally) were checked. Manual inspection assessed presence of distinct group for individual transitions above noise level with matching peak shapes and relative ratio of individual transition signals matching one observed for synthetic peptide. Manual inspection showed that 62% of chromatogram groups lacked peptide signal. Skyline peak detection exhibited certain instability depending on exact set of runs imported and also randomly with rescoring attempts, probably due to lack of internal standards.

In 16% of groups peak was detected and no adjustment to integration limits established by Skyline was required, in 22% of groups some adjustment of integration limits was needed. False negative results, where Skyline missed a peak group that was found during manual inspection, were not observed. In 30% of groups both Skyline and manual inspection reported no peak detection. In 31% of groups detection was reported (at least one transition area above zero for a peptide in a run) by Skyline, but rejected after manual inspection (45% false positive rate).

Typically Skyline is used to process established quantitative methods with internal standards. In that case, high-confidence peak of internal standard is detected, limits of integration are determined by it and cross-product (dotp) between natural peptide and internal standard transition signals is used to assess peak detection quality with low-scoring peaks being rejected. In addition, results that fall below preliminarily determined limits of detection are rejected. In this work detection itself was studied, internal standards were not used and limits of detection were not determined. Manual inspection of a single run for 123 peptides takes about 5 minutes for an experienced analyst, but inspection of 96 runs would take 8 hours. Both long inspection time and distribution of manual inspection between several analysts might lead to biases in peak detection and should be avoided. Thus, in order to exclude false positive peptide detections, properties of manually inspected were explored and used to filter out false positive detections that were typically low intensity or noise level, had different retention time than synthetic peptide under identical separation conditions or had different relative contribution of individual transitions into total peptide signal, which is expected due to stability of fragmentation pattern.

Since several LC-MS replicates of a synthetic peptide pool were run during analysis of each plasma batch, median retention time of peptide and normalized vectorized representation for relative transition signals (as a unit vector in -dimensional Euclidean space for peptide with transitions) were computed for low number of standard sample runs with unambiguous high-confidence peaks inspected manually. Retention time shift, cross-product of vectorized relative intensities (i.e. cosine of angle between vectors representing relative intensities), total ion current intensity and individual transition intensities were studied. Based on two reports, one produced by Skyline after automatic built-in peak detection algorithm and one produced after manual inspection of peaks in Skyline, comparison of total ion current, fraction of transitions with area falling below a defined threshold, absolute retention time shift from median value observed in pooled standard sample and cross-product (dotp) for three groups of signals reported with built-in Skyline peak detection algorithm - those where peak detection was rejected after manual inspection (false positives), where peak limits were adjusted (true positives) and those where no change to algorithms’ choice was made (true positives) (Figures 1-4). Peaks reported by Skyline were assessed based on four criteria - total ion current of above 10^5^, retention time difference not more than 0.1 minute from median time determined for standard runs, cross-product of relative transition intensities computed against standard runs 0.97 or above and at least 80% of transitions have area at least 10^4^. Peaks that matched at least 3 criteria were marked as detected. Comparison to manually assessed plasma sample runs showed 97% retention of true peak detections and 5% false discovery (peak detection) rate.

**Fig. 1.**
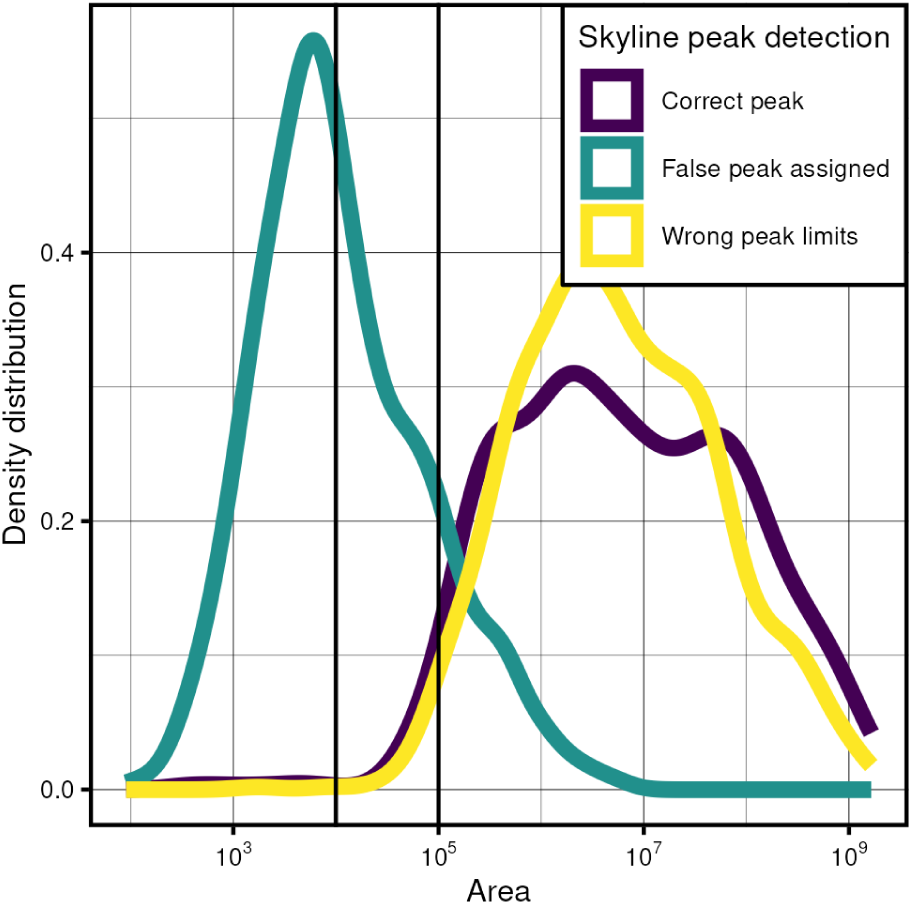
Area distribution

**Figure 2.**
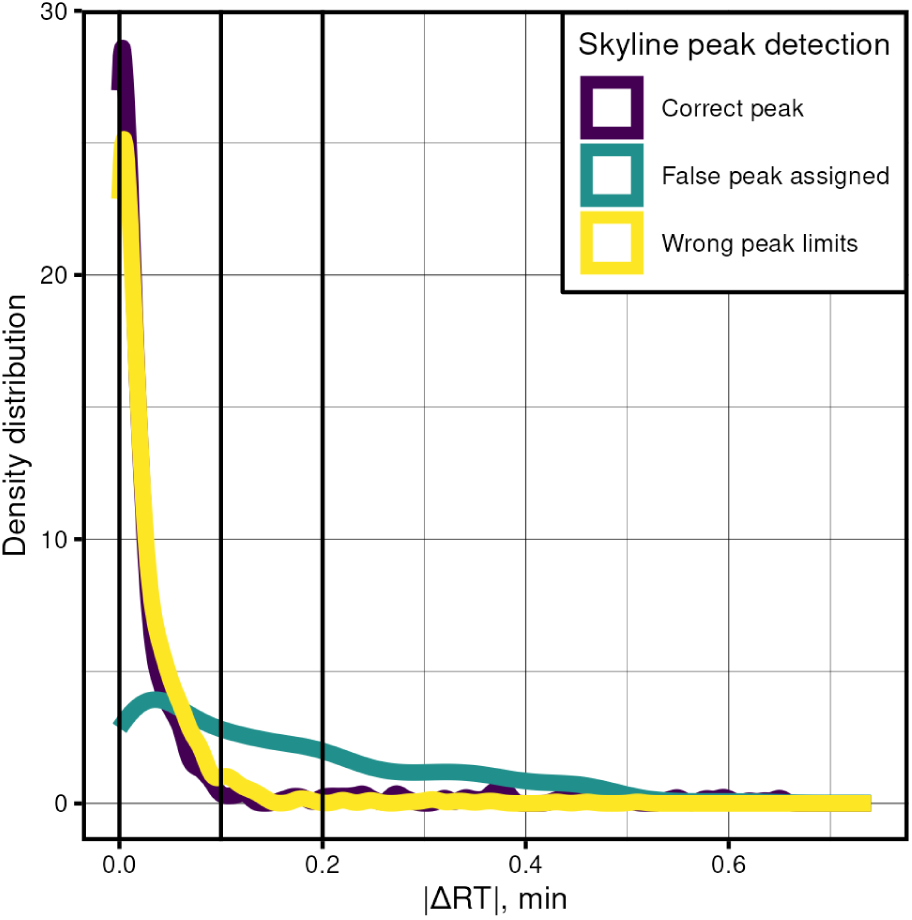
Difference in retention time distribution

**Figure 3.**
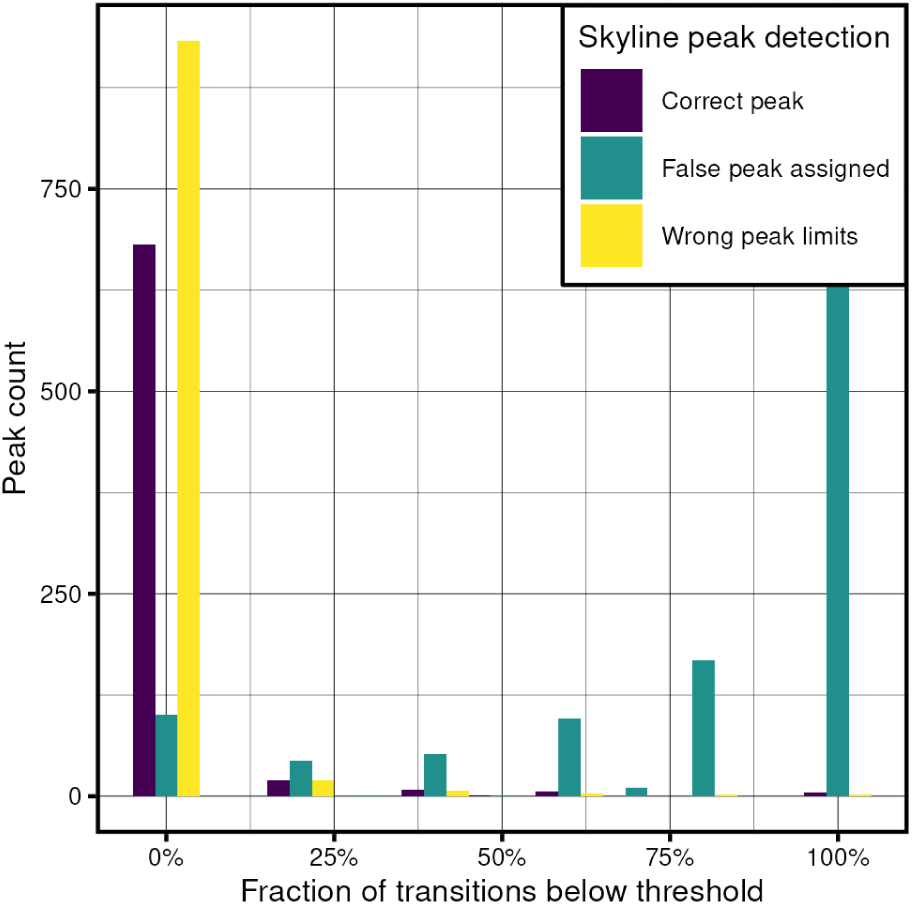
Fraction of transitions below threshold

**Figure 4.**
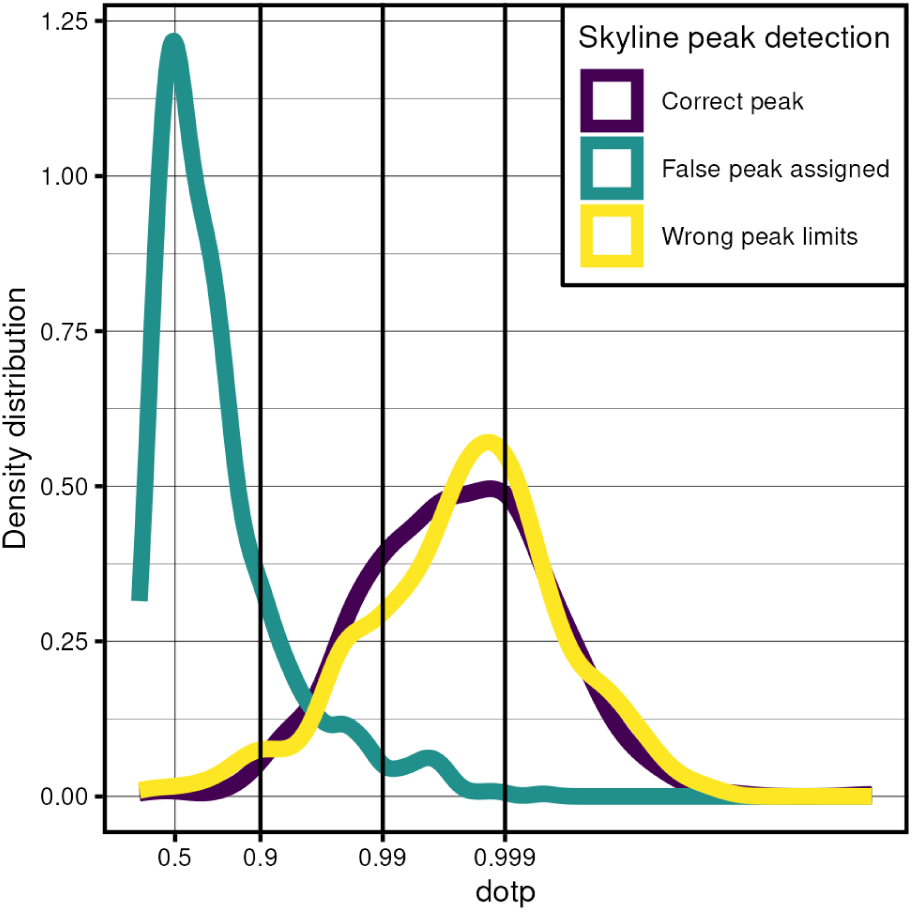
Cross product of relative transition intensities

#### Detection rate

Detection rate of 123 peptides of albumin, ceruloplasmin and C-reactive protein was computed for a collection of 96 healthy volunteers samples (Figure 5). Detection rates covered whole range from 0 to 1 with 63 peptides being never detected and median detection rate for the rest of peptides equal to 0.74. For C-reactive protein single peptide (ESDTSYVSLK) was observed in a collection of healthy volunteers plasma.

**Figure 5.**
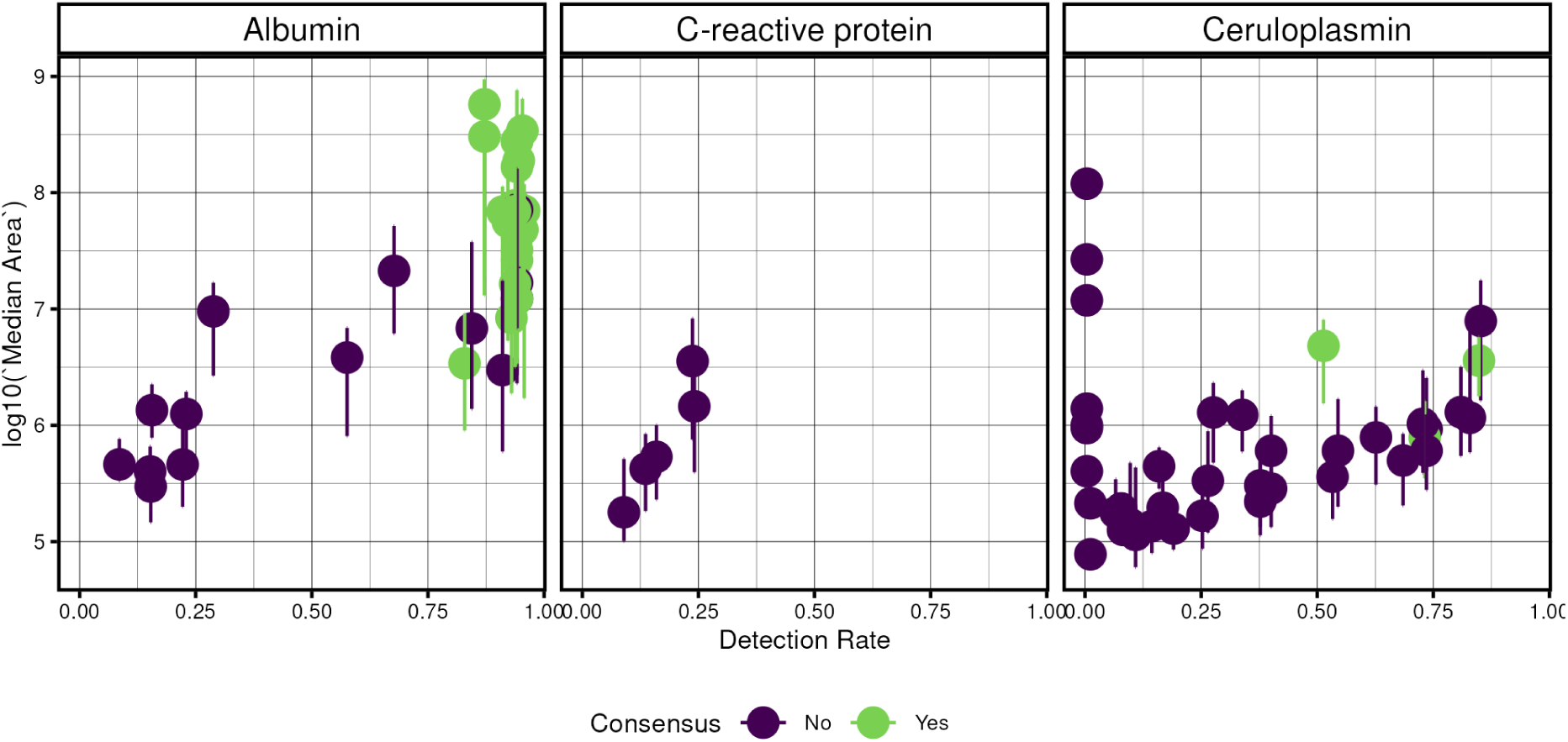
Peptide detection rates vs median peptide TIC Lines represent 10th and 90th percentiles.

Since albumin and ceruloplasmin are high-abundance proteins, residual non-alkylated cysteine-containg peptides might be expected to be observed. Cysteine-containing peptides in unmodified form were not detected in plasma samples except for 3 of 5 peptides with highest signal of alkylated form. Two of three visible peptides contained two vicinal (consecutive) cysteines (<u>10.1016</u>/j.<u>jmb.2017.03.017</u>), of which one peptide in 73% of samples (2 Da mass shift is within transmission window for +2 and +3 charged ions) and the other bearing N-terminal cysteine was observed in 1% of samples (probably low due to additional in vitro carbamylation by urea during processing, <u>10.1006/abio.1998.2970</u>).

Unmodified cysteine-bearing peptides were excluded from further analysis. At the same time, alkylated cysteine-containing peptides were among most intense most frequently observed. While these peptides are typically not used when peptide-level internal standards are used due to necessity of separate alkylation procedure for them, QConCAT/QPreST-based quantitation might be performed with cysteine-containing peptides. All observed peptides could be attributed to canonical isoforms of proteins.

Detection rate had high correlation with median log-transformed observed peptide total ion current, i.e. sum of several MRM transitions (Figure 5, Pearson’s ρ 0.74). This might point to conclusion that peptide’s proteotypicity is linked to limit of detection achieved for it, but both albumin and ceruloplasmin are amongst most abundant proteins in blood plasma and detected peptides exhibit reproducible high-intensity signal. This contradiction might be explained by interplay of genetical variation, allele frequency and differential allele-specific expression (<u>10.1016</u>/<u>j.t</u>ig<u>.2014.03.003</u>) or varying level of post-translational modifications.

Human Plasma 2025-08 PeptideAtlas Build combines results from 30695 DDA LC-MS runs acquired within 356 experiments performed independently using a variety of sample preparation approaches and LC-MS configurations. It summarizes each peptides’ detection across all experiments with Empirical Proteotypic Score (By PeptideAtlas definition: In PeptideAtlas, the fraction of observations of this protein that are supported by at least one observation of this peptide ). Due to albumin and ceruloplasmin being major proteins of plasma with important physiological functions (albumin maintains oncotic pressure transports a variety of water-insoluble substances, ceruloplasmin transports copper and exhibits ferroxidase activity), detection rate determined for them on a collection of healthy individuals can be treated as peptide proteotypicity. In fact, since several albumin peptides were detected in every sample, detection rate for albumin peptides is proteotypicity by definition (rate of peptide detection whenever protein was detected). As for ceruloplasmin, while none of it’s peptides reached 100% detection rate, it is unlikely that any plasma sample did not contain ceruloplasmin (and all samples contained blood plasma due to albumin being detected in every sample).

Observed detection rates were compared to from Human

Plasma 2025-08 PeptideAtlas Build. Albumin peptides exhibited mostly higher detection rates than reported in PeptideAtlas, and ceruloplasmin peptides - lower ones (Figure 6). A possible explanation of these discrepancies and their opposing direction is an interplay of methodological and populational variance. had low correlation with median log-transformed observed peptide total ion current (Pearson’s ρ 0.06), which means that high proteotypicity score is a poor predictor for high signal intensity and to some extent, low limit of detection.

**Figure 6.**
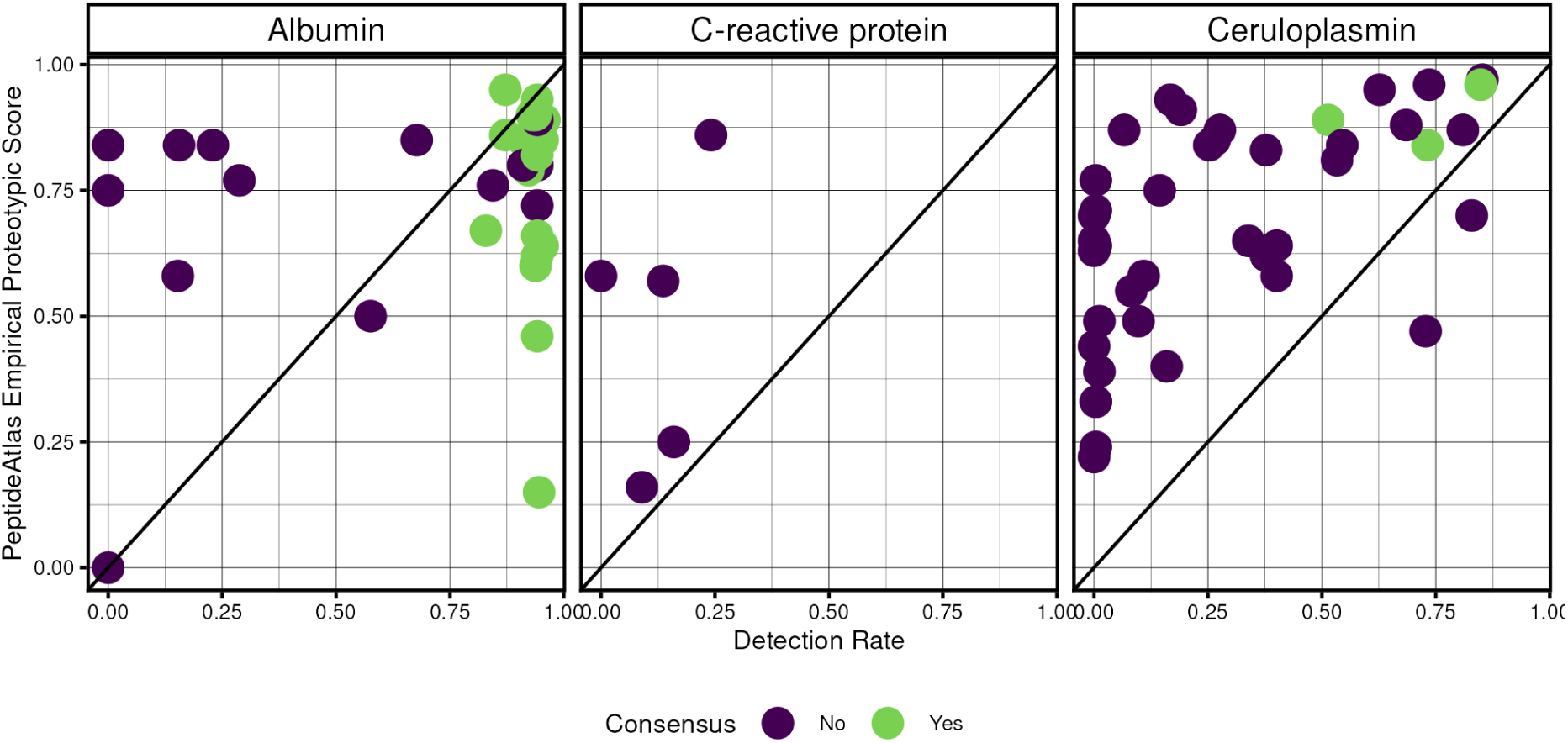
Peptide detection rates vs PeptideAtlas Empirical Proteotypicity Score

Besides proteotypicity, quantotypicity (<u>10.1089/omi.2011.0156</u>) of peptides is important for the development of targeted quantitation methods. Though label-free quantitation requires sample normalization under the assumption that among multiple quantified proteins few are differentially abundant and targeted quantification is typically performed with isotopically-labelled internal peptide standards, synthesis of 50-100 isotopically labelled standards was deemed unreasonable. Pearson’s correlation coefficient of TIC was computed for all peptide pairs observed together in at least 41 of 96 samples. For pairs of peptides originating from different proteins only 2% had Pearson’s ρ was 0.9 or higher and for pairs of peptides originating from the same protein 25% had correlation coefficient that high (Figure 7). For every protein a peptide graph was built with edges between peptides having correlation coefficient of 0.9 or above. Peptides belonging to the largest connected component of such graph were called “consensus” peptides. The above mentioned difference in correlation between intra-protein or cross-protein peptide pairs holds only when both peptides belong to these consensus peptides. “Consensus” peptides tend to be more frequently detected and have higher intensity (figures 5 and 6). Since their response is mutually concordant, they might be the best candidates for quantotypicity testing.

**Figure 7.**
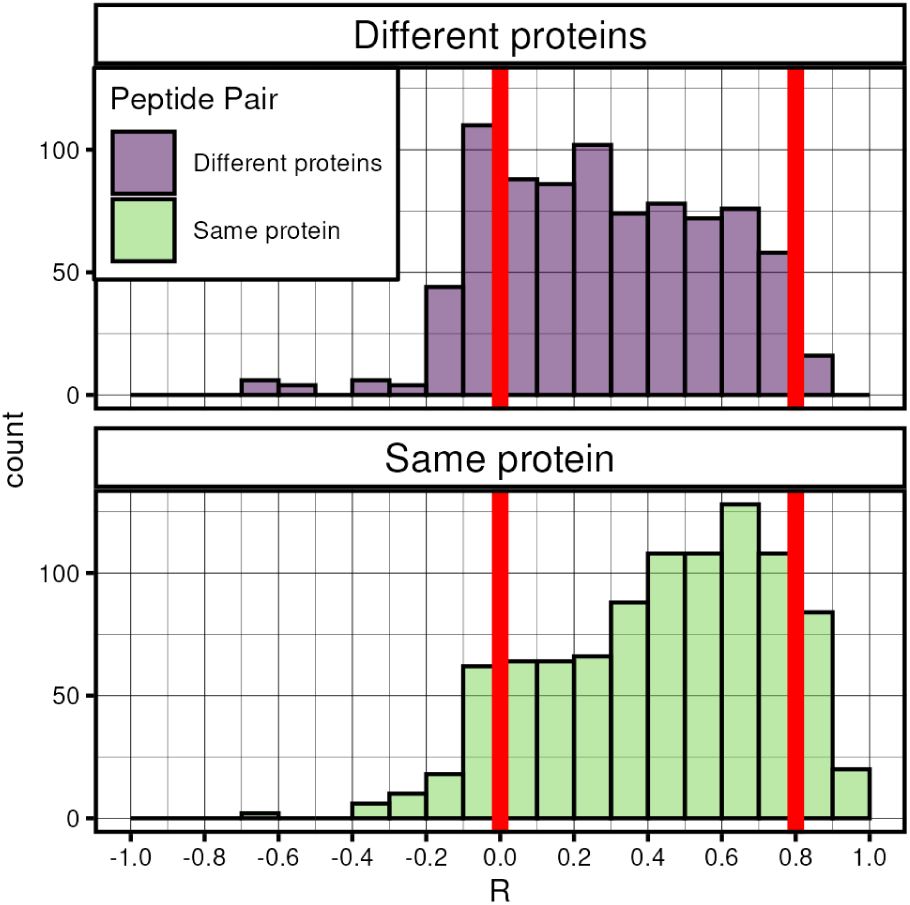
Peptide TIC correlation

#### Biological and technical factors in proteotypicity

When established targeted detection for high-abundant proteins is considered, one can expect event of choosing a sample where genetic and other slowly-changing biological factors come in favor of producing targeted peptide from targeted protein and event of detecting a peptide in obtained sample (influence of technical factors) to be independent. In that case, probability of detecting a peptide in a single sample is a product of two probabilities - the probability of choosing a suitable sample and probability of successfully detecting a peptide in a sample where peptide exists. If no systematic error or bias occurs during sample processing and data acquisition, one can consider consecutive tests to be independent and probabilities of detection to be identically distributed and in that case is described by a Bernoulli process. Probability of k successes in n trials with probability of success in a single trial being p is described by a binomial distribution.

Detection rates in daily replicates of ICU patients’ plasma were used to estimate (with MLE) contribution of sample processing (technical or rapidly changing biological factors) factors in limited proteotypicity. Some peptides were not observed for certain (but not all) patients at all. Probability estimates in range from 0.01 to 0.95 (median 0.6) were obtained for 70 peptides. From detection rates and estimates of processing-related probabilities, contribution of genetic factors was estimated by division of apparent detection rate by processing-related probability. While whole range of probabilities was covered, for vast majority of peptides biology-related contribution to limited proteotypicity is negligible for most peptides studied (Figure 8).

**Figure 8.**
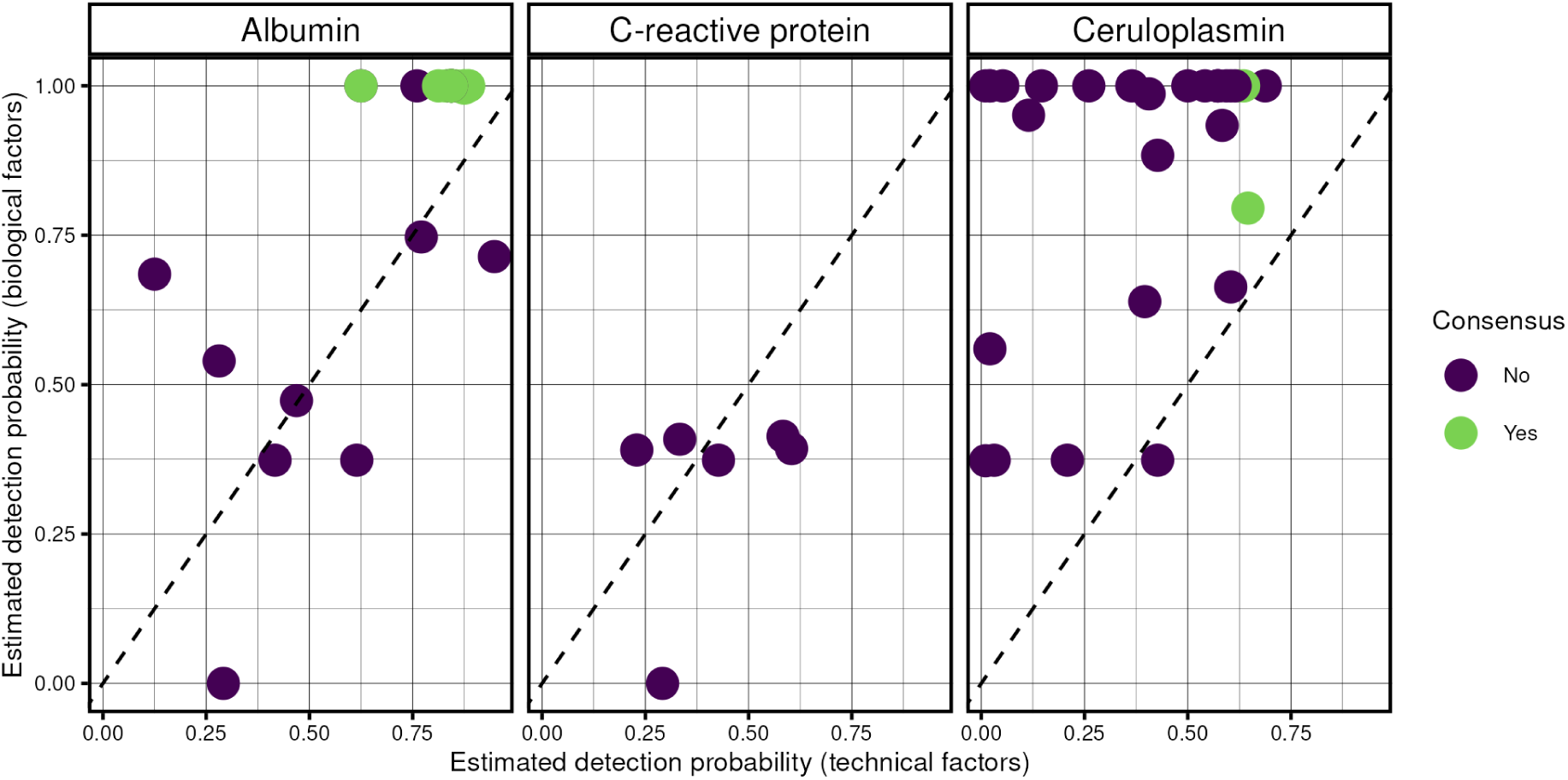

## Conclusion

Peptide proteotypicity is a long-known concept of utmost importance for practical applications of bottom-up proteomics and many of it’s properties are known. Multiple prediction approaches and summarized common empirical experience are available for development of targeted proteomics methods. We attempted to assess and compare proteotypicity of peptides in a custom in-house developed targeted bottom-up proteomics analysis of human plasma. We show that direct in-house empirical assessment of peptide proteotypicity is possible, it’s results do not fully match with common predictions or observations. Discrepancies might be explained by either known platform-dependency of proteotypicity or biological effects. Definite answer requires further research most probably for each separate protein target and peptide. Lack of reproducible detection on a population scale might be attributed to peptide properties or populational variation in either sequence (SAPs) or protein abundance. The first one can and should be tested with recombinant protein spike-ins an if it is shown not to be the case, genetic variation, abundance or PTM hypotheses should be explicitly tested.

Targeted bottom-up proteomics-based tests’ sensitivity (i.e. probability of positive result in a population of patients with a condition to be diagnosed) can be compromised by low proteotypicity (i.e. probability of peptide detection in a set of samples with a protein to be detected) and lead to high false negative rate and low negative predictive value. We propose that proteotypicity of peptide should be defined for a specific sample (for example, human blood plasma) processed on a specific platform, including sample preparation and detection technologies. Targeted bottom-up quantitation methods to be used for biomarker discovery or clinical analysis have to be characterized in term of proteotypicity of peptides (or peptide sets) used for each protein detection in a certain population. Technical and biological factors influencing proteotypicity can be estimated. Limited proteotypicity should be accounted for during interpretation of test or biomarker figures of merit and increase in sensitivity might be expected after translation from bottom-up to top-down approaches.

## Abbreviations

LC: liquid chromatography
MS: mass-spectrometry
LC-MS: liquid chromatography coupled to mass-spectrometry
MRM: multiple reaction monitoring
SRM: single reaction monitoring
DDA: data-dependent acquisition
IDA: information-dependent acquisition
AMT: accurate mass and time
DIA: data-independent acquisition
SWATH: sequential window acquisition of all theoretical fragment ions
MS1: parent ion spectrum
MS2: product ion spectrum
TOF: time-of-flight
ESI: electrospray ionization
HPLC: high-performance liquid chromatography
UHPLC: ultra-high performance liquid chromatography
LOD: limit of detection
AAA: amino acid analysis
MIDAS: multiple reaction monitoring-initiated detection and sequencing
SVM: support vector machine
iRT: indexed retention time
DP: declustering potential
CE: collision energy
PTM: post-translational modification
ICU: Intensive Care Unit

