## Supplementary Material for "Direct empirical in-house assessment of peptide proteotypicity for targeted proteomics"

### Supplementary Material

#### Code for peptide property assessment

##### Script to access neXtProt

```
# Script to collect and store information about proteins from neXtProt via
REST API and tidyverse
# This script modifies neXtProt responses to allow caching them for a year!

library(magrittr)
library(httr2)
library(dplyr)
library(readr)
library(tidyr)
library(stringi)
library(Peptides)
library(pbapply)
library(parallel)
library(data.table)
library(ggplot2)

# library(dtplyr)

options(readr.show_col_types = FALSE)

# Detect database and API releases, set up cache and API call wrapper

versions <-
  'https://api.nextprot.org/' %>%
  request %>%
  req_url_path_append('release-info.json') %>%
  req_perform %>%
  resp_body_json

cache_directory <-
  file.path('data/nextprot_rest_api_cache',
            paste0(versions$versions, collapse = '_'))

if (!dir.exists(cache_directory))
  dir.create(cache_directory)
```

```
cache_update_and_force_Expire <-
  function(resp,
           Expires = format(
             strptime(resp_header(resp, 'Date'),
                      format = '%a, %d %b %Y %H:%M:%S',
                      tz = 'GMT') + 60 * 60 * 24 * 365,
             '%a, %d %b %Y %H:%M:%S %Z'
           ),
           verbose = FALSE) {
    cache_path <- httr2:::req_cache_path(resp$request)
    cached_time <- NULL
    cached_Expires <- NULL

    if (file.exists(cache_path)) {
      if (verbose)
        message('Has cache!')
      cached <- readRDS(cache_path)
      cached_time <-
        strptime(resp_header(cached, 'Date'),
                 format = '%a, %d %b %Y %H:%M:%S',
                 tz = 'GMT')
      cached_Expires <- resp_header(cached, 'Expires')
    }

    resp_time <-
      strptime(resp_header(resp, 'Date'),
               format = '%a, %d %b %Y %H:%M:%S',
               tz = 'GMT')

    if (is.null(cached_time) || (!is.null(cached_time) &&
                                 ((resp_time > cached_time) ||
                                  (!is.null(cached_Expires) &&
                                   cached_Expires != Expires)
                                 ))) {
      if (verbose &&
          !is.null(cached_time) &&
          resp_time > cached_time)
        message('Stale cache!')
      if (verbose &&
          !is.null(cached_Expires) &&
          cached_Expires != Expires)
        message('Expires mismatch!')
      if (verbose)
        message('Update Expire header and cache!')
      resp[['headers']][['Expires']] <- Expires
      saveRDS(resp, cache_path)
```

```
    }

    resp
  }

req <-
  'https://api.nextprot.org/' %>% request %>% req_cache(
    path = cache_directory,
    max_size = 1024 * 1024 * 1024 * 10,
    use_on_error = TRUE,
    debug = FALSE
  )

get_nextprot <-
  function(x,
           verbosity = 0L,
           verbose_cache = FALSE,
           pause = NULL) {
    if (!is.null(pause))
      Sys.sleep(pause)
    req %>%
      req_url_path_append(x) %>%
      req_perform(verbosity = verbosity) %>%
      cache_update_and_force_Expire(verbose = verbose_cache)
  }

cluster <- makeForkCluster(nnodes = 64L)

# Get database and API release
versions <- get_nextprot('release-info.json')

# Get chromosome names, chromosome summary and all entry accessions
chromosomes <-
  'chromosomes.json' %>%
  get_nextprot %>%
  resp_body_json %>%
  unlist
chromosome_summary <-
  'chromosome-reports/summary.json' %>%
  get_nextprot %>%
  resp_body_json %>%
  lapply(as_tibble) %>%
  bind_rows(.id = 'chromosome')

accessions <-
  'entry-accessions.json' %>%
```

```r
  get_nextprot %>%
  resp_body_json %>%
  unlist

# Get genome locations
chromosome_export <-
  chromosomes %>% paste0('.tsv') %>%
  paste('chromosome-report', 'export', ., sep = '/') %>%
  lapply(get_nextprot) %>%
  lapply(resp_body_string) %>%
  lapply(function(x) {
    read_tsv(x, col_types = 'ccciicc')
  }) %>% bind_rows
chromosome_export %<>%
  mutate(`UniProt Entry` = sub('^NX_', '', `neXtProt AC`))

# Get isoform sequences

isoform_sequences <-
  'isoforms.tsv' %>%
  get_nextprot %>%
  resp_body_string %>%
  read_tsv

isoform_sequences %<>%
  mutate(`neXtProt AC` = sub('-[[:digit:]]+$', '', isoform)) %>%
  relocate(c('isoform', 'neXtProt AC', 'md5', 'sequence'))

isoform_sequences %>%
  group_by(`neXtProt AC`) %>%
  summarise(`Isoforms per Entry` = n_distinct(isoform)) %>%
  group_by(`Isoforms per Entry`) %>%
  summarise(Entries = n_distinct(`neXtProt AC`))

# Get RefSeq mapping

refseq_map <-
  c('mapping', 'nextprot_refseq.tsv') %>%
  get_nextprot %>%
  resp_body_string %>%
  read_tsv(col_names = c('neXtProt AC', 'RefSeq Protein', 'RefSeq mRNA'))


# Get PTM information

if (file.exists(file.path(cache_directory, 'ptms.rds'))) {
```

```
  ptms <- readRDS(file.path(cache_directory, 'ptms.rds'))
} else {
  ptms <-
    accessions %>% paste('entry', ., 'ptm.json', sep = '/') %>% pblapply(cl =
cluster, get_nextprot) %>% pblapply(cl = cluster, resp_body_json)

  saveRDS(ptms, file.path(cache_directory, 'ptms.rds'))
}

if (file.exists(file.path(cache_directory, 'ptms_export.rds'))) {
  ptms_export <-
    readRDS(file.path(cache_directory, 'ptms_export.rds'))
} else {
  ptms_export <- pblapply(cl = cluster, ptms, function(x) {
    e <-
      tibble(
        `neXtProt AC` = x$entry$uniqueName,
        `isoform Count` = length(x$entry$isoforms)
      )
    p <- NULL
    if (length(x$entry$annotationsByCategory) > 0L)  {
      p <- lapply(x$entry$annotationsByCategory[[1L]], function(z) {
        sel <- sapply(z, class)
        sel <-
          names(sel[sel %in% c('integer', 'numeric', 'logical', 'character')])
        d <-
          as_tibble(z[sel]) %>% bind_rows
        l <-
          lapply(z$targetingIsoformsMap, function(y) {
            sel <- sapply(y, class)
            sel <-
              names(sel[sel %in% c('integer', 'numeric', 'logical',
'character')])
            as_tibble(y[sel])
          }) %>% bind_rows
        cross_join(d, l, copy = TRUE)
      }) %>% bind_rows
      e <- cross_join(e, p, copy = TRUE)
    }
    e
  }) %>% bind_rows

  ptms_export %<>% filter(!is.na(cvTermAccessionCode)) %>% select(
    `neXtProt AC`,
    isoformName,
    firstPosition,
```

```
    lastPosition,
    cvTermAccessionCode,
    cvTermName,
    cvTermDescription
  ) %>% pivot_longer(cols = c(firstPosition, lastPosition)) %>% select(-name)
%>% unique %>% rename(position = value)

  saveRDS(ptms_export, file.path(cache_directory, 'ptms_export.rds'))
}

ptm_map <-
  ptms_export %>% left_join(isoform_sequences %>% select(isoform, md5),
                            by = c('isoformName' = 'isoform')) %>%
group_by(md5, position) %>% summarise(cvTermName = paste0(cvTermName, collapse
= ';'))

# Get sequence variants information

if (file.exists(file.path(cache_directory, 'variants_export.rds'))) {
  variants_export <-
    readRDS(file.path(cache_directory, 'variants_export.rds'))
} else {
  if (file.exists(file.path(cache_directory, 'variants.rds'))) {
    variants <-
      readRDS(file.path(cache_directory, 'variants.rds'))
  } else {
    variants <-
      accessions %>% paste('entry', ., 'variant.json', sep = '/') %>%
pblapply(cl = cluster, get_nextprot) %>% pblapply(cl = cluster,
resp_body_json)

    saveRDS(variants, file.path(cache_directory, 'variants.rds'))
  }

  variants_export <-
    pblapply(cl = cluster, variants[1L:128L], function(x) {
      e <-
        tibble(
          `neXtProt AC` = x$entry$uniqueName,
          `isoform Count` = length(x$entry$isoforms)
        )
      p <- NULL
      if (length(x$entry$annotationsByCategory) > 0L)  {
        p <- lapply(x$entry$annotationsByCategory[[1L]], function(z) {
          sel <- sapply(z, class)
          sel <-
```

```
            names(sel[sel %in% c('integer', 'numeric', 'logical',
'character')])
          d <-
            as_tibble(z[sel]) %>% bind_rows
          l <-
            lapply(z$targetingIsoformsMap, function(y) {
              sel <- sapply(y, class)
              sel <-
                names(sel[sel %in% c('integer', 'numeric', 'logical',
'character')])
              as_tibble(y[sel])
            }) %>% bind_rows
          sel <- sapply(z$variant, class)
          sel <-
            names(sel[sel %in% c('integer', 'numeric', 'logical',
'character')])
          v <- as_tibble(z$variant[sel])
          r <- cross_join(d, v, copy = TRUE)
          r <- cross_join(r, l, copy = TRUE)
          r
        }) %>% bind_rows
        e <- cross_join(e, p, copy = TRUE)
      }
      e
    }) %>% bind_rows
}

variants_export %<>% filter(!is.na(firstPosition)) %>% rowwise %>%
mutate(affected = list(firstPosition:lastPosition)) %>% select(`neXtProt AC`,
isoformAccession, affected, original, variant) %>% unnest_longer(col =
affected, values_to = 'Position')

variants_map <-
  variants_export %>% left_join(isoform_sequences %>% select(isoform, md5),
                                by = c('isoformAccession' = 'isoform')) %>%
group_by(md5, Position) %>% summarise(
                                  original = paste0(unique(original), collapse
= ';'),
                                  variant = paste0(unique(variant), collapse =
';')
                                )

# List different protein chains

chains <-
  isoform_sequences %>% group_by(md5, sequence) %>%
```

```
  summarise(isoform = list(sort(unique(isoform))), `neXtProt AC` =
list(sort(unique(`neXtProt AC`)))) %>%
  rowwise %>%
  mutate(
    `Protein Length` = nchar(sequence),
    `isoform Count` = length(isoform),
    `neXtProt AC Count` = length(`neXtProt AC`)
  )

# Annotate each residue in chains

exploded <-
  chains %>%
  select(md5, sequence) %>%
  distinct %>%
  group_by(md5) %>%
  mutate(Length = nchar(sequence)) %>%
  mutate(Residues = {
    x <-
      strsplit(sequence, split = '|')
    names(x) <-
      1L:length(x)
    x
  }, .keep = 'none') %>%
  unnest_longer(col = Residues,
                indices_to = 'Position',
                values_to = 'Residue') %>%
  relocate(md5, Position, Residue) %>%
  mutate(`Cleave Before` = replace(
    Residue != 'P' &
      (Position == 1L |
         lag(Residue) == 'K' |
         lag(Residue) == 'R'),
    Position == 1L,
    FALSE
  )) %>%
  mutate(`Peptide N` = cumsum(`Cleave Before`) + 1L) %>%
  mutate(`Cleave After` = lead(`Cleave Before`, default = FALSE))

exploded %<>% left_join(ptm_map, by = c('md5' = 'md5', 'Position' =
'position'))
exploded %<>% left_join(variants_map, by = c('md5' = 'md5', 'Position' =
'Position'))

sap_changes_sites_1 <-
  exploded %>% filter(`Cleave After`, !is.na(variant)) %>% select(md5,
```

```
`Peptide N`) %>% mutate(`Peptide N` = `Peptide N` + 1L,

`Affected by Variant in Previous Peptide` = TRUE)
sap_changes_sites_2 <-
  exploded %>% filter(`Cleave Before`, !is.na(variant), grepl('P', variant))
%>% select(md5, `Peptide N`) %>% mutate(`Peptide N` = `Peptide N` - 1L,

`Affected by Variant in Next Peptide` = TRUE)

exploded %<>% left_join(sap_changes_sites_1, by = c('md5', 'Peptide N')) %>%
left_join(sap_changes_sites_2, by = c('md5', 'Peptide N'))

# Compile tryptic peptide sequences

cleaved <-
  exploded %>%
  group_by(md5, `Peptide N`) %>%
  summarise(
    `Peptide Sequence` = paste0(Residue, collapse = ''),
    `Peptide Start` = first(Position),
    `Peptide End` = last(Position),
    `N-terminal Peptide` = !any(`Cleave Before`),
    `C-terminal Peptide` = !any(`Cleave After`),
    `Phosphorylated Residues` = sum(
      cvTermName %in% c('Phosphoserine', 'Phosphothreonine',
'Phosphotyrosine'),
      na.rm = TRUE
    ),
    `Total Modified Residues` = sum(!is.na(cvTermName)),
    `Sequence Variants` = any(
      !is.na(variant),
      !is.na(`Affected by Variant in Next Peptide`),
      !is.na(`Affected by Variant in Previous Peptide`)
    )
  ) %>% ungroup %>% mutate(`Peptide Length` = `Peptide End` - `Peptide Start`
+ 1L) %>%
  group_by(md5) %>%
  mutate(`Next Site` = lead(`Peptide Length`, default = 100000L)) %>%
  mutate(`Previous Site` = lag(`Peptide Length`, default = 100000L))

# List unique peptide sequences and compute their properties

peptides <-
  cleaved %>% ungroup %>% select(`Peptide Sequence`, `Peptide Length`) %>%
distinct
```

```
peptides %<>% mutate(
  `Has C` = stri_count_fixed(`Peptide Sequence`, 'C'),
  `Has ^Q` = stri_count_regex(`Peptide Sequence`, '^Q'),
  `Has ^E` = stri_count_fixed(`Peptide Sequence`, '^E'),
  `Has N` = stri_count_fixed(`Peptide Sequence`, 'N'),
  `Has Q` = stri_count_fixed(`Peptide Sequence`, 'Q'),
  `Has X` = stri_count_fixed(`Peptide Sequence`, 'X'),
  `Has O` = stri_count_fixed(`Peptide Sequence`, 'O'),
  `Has U` = stri_count_fixed(`Peptide Sequence`, 'U'),
  `Has M` = stri_count_fixed(`Peptide Sequence`, 'M'),
  `Has W` = stri_count_fixed(`Peptide Sequence`, 'W'),
  `Has DP` = stri_count_fixed(`Peptide Sequence`, 'DP'),
  `Has NP` = stri_count_fixed(`Peptide Sequence`, 'NP'),
  `Has NG` = stri_count_fixed(`Peptide Sequence`, 'NG'),
  `Has KP` = stri_count_fixed(`Peptide Sequence`, 'KP'),
  `Has RP` = stri_count_fixed(`Peptide Sequence`, 'RP')
)

peptides %<>% mutate(
  `MW` = mw(
    `Peptide Sequence`,
    monoisotopic = TRUE,
    aaShift = c(C = 57.021464)
  ),
  `m/z +2` = mz(`Peptide Sequence`, charge = 2L, cysteins = 57.021464),
  `m/z +3` = mz(`Peptide Sequence`, charge = 3L, cysteins = 57.021464),
  `Hydrophobicity` = hydrophobicity(`Peptide Sequence`)
)

# Compute peptide penalties based on peptide's position in a chain, PTMs or
Sequence Variants

cleaved %<>% mutate(
  `Position Penalty` =  `N-terminal Peptide` * 1000L +
    `C-terminal Peptide` * 1000L +
    (`Next Site` < 5L) +
    (`Previous Site` < 5L) +
    (`Next Site` < 3L) * 1000L +
    (`Previous Site` < 3L) * 1000L,
  `Modification Penalty` = `Total Modified Residues` * 1000L,
  `Variant Penalty` = `Sequence Variants` * 1000L
)

# Compute peptide penalty based on it's amino acid composition

peptides %<>% mutate(
```

```
  `Composition Penalty` =
    `Has C` * 1000L + # Prohibited (standards should be modified)
    `Has W` * 1000L + # Prohibited (oxidizes heavily with several channels)
    `Has N` + # Deamidation
    `Has Q` + # Deamidation
    `Has M` + # Oxidation
    `Has ^E` * 1000L + # These N and Q are penalized additionaly, they are
prone to cyclization during standard synthesis
    `Has ^Q` * 1000L + # These N and Q are penalized additionaly, they are
prone to cyclization during standard synthesis
    `Has NG` + # These N and Q are penalized additionaly, they are more prone
to modification
    `Has NP` + # These N and Q are penalized additionaly, they are more prone
to modification
    `Has DP` + # Unstable
    `Has KP` + # This cleavage occurs but slowly
    `Has RP` + # This cleavage occurs but slowly
    `Has O` * 1000L + # Prohibited (synthesis / biosynthesis problems)
    `Has X` * 1000L + # Prohibited (synthesis / biosynthesis problems)
    `Has U` * 1000L + # Prohibited (synthesis / biosynthesis problems)
    (`Peptide Length` < 5L) * 1000L + # Short are less specific and too
hydrophylic
    (`Peptide Length` > 30) * 1000L + # Long are less stable and more
expensive
    (`m/z +2` < 300 &
       `m/z +3` < 300) * 1000L + # Below 300 Da the background is high and MS
less stable
    (`m/z +2` > 1500 &
       `m/z +3` > 1500) * 1000L + # Above 1500 Da MS less stable
    (Hydrophobicity < -2) + (Hydrophobicity > 1) # Early and late peptides
depend too much on stationary phase properties
)

peptide_sources <-
  chains %>% group_by(md5) %>%
  select(md5, isoform, `neXtProt AC`) %>%
  right_join(
    cleaved %>% ungroup %>% select(`Peptide Sequence`, md5) %>% distinct,
    by = 'md5',
    keep = NULL,
    relationship = 'one-to-many',
    unmatched = 'error',
    multiple = 'all'
  ) %>%
  group_by(`Peptide Sequence`) %>%
  summarise(
```

```
    md5 = list(sort(unique(md5))),
    isoform = list(sort(unique(
      unlist(isoform, use.names = FALSE)
    ))),
    `neXtProt AC` = list(sort(unique(
      unlist(`neXtProt AC`, use.names = FALSE)
    )))
  ) %>%
  rowwise %>%
  mutate(
    `md5 Count` = length(md5),
    `isoform Count` = length(isoform),
    `neXtProt AC Count` = length(`neXtProt AC`)
  )


{
  hbpp255_peptides <-
    read_lines('data/raw_files/peptide_lists/hbpp255.txt')

  table(hbpp255_peptides %in% peptide_sources$`Peptide Sequence`)
  peptide_sources %>% filter(`Peptide Sequence` %in% hbpp255_peptides) %>%
group_by(`neXtProt AC Count`) %>% summarise(Peptides = n())

  borchers_peptides <-
    read_lines('data/raw_files/peptide_lists/borchers.txt')

  table(borchers_peptides %in% peptide_sources$`Peptide Sequence`)
  peptide_sources %>% filter(`Peptide Sequence` %in% borchers_peptides) %>%
group_by(`neXtProt AC Count`) %>% summarise(Peptides = n())

  synthesized_peptides <-
    read_lines('data/raw_files/peptide_lists/synth_summary_20240226.txt')

  table(synthesized_peptides %in% peptide_sources$`Peptide Sequence`)
  peptide_sources %>% filter(`Peptide Sequence` %in% synthesized_peptides) %>%
group_by(`neXtProt AC Count`) %>% summarise(Peptides = n())

  enc_proteins <- read_lines('data/raw_files/protein_lists/enc.txt')
  setdiff(enc_proteins, chromosome_export$`UniProt Entry`)
  enc_AC <-
    chromosome_export %>% filter(`UniProt Entry` %in% enc_proteins) %>%
select(`neXtProt AC`) %>% unlist(use.names = FALSE)

  hbpp255_proteins <-
    read_lines('data/raw_files/protein_lists/hbpp255.txt') %>% strsplit(split
```

```
= '|', fixed = TRUE) %>% sapply(`[[`, 2L) %>% strsplit(split = '-', fixed =
TRUE) %>% sapply(`[[`, 1L) %>% sort %>% unique %>% setdiff(c('P00760',
'P02769'))
  setdiff(hbpp255_proteins, chromosome_export$`UniProt Entry`)
  hbpp255_AC <-
    chromosome_export %>% filter(`UniProt Entry` %in% hbpp255_proteins) %>%
select(`neXtProt AC`) %>% unlist(use.names = FALSE)

  covid31_proteins <-
    read_lines('data/raw_files/protein_lists/covid31.txt') %>% sort %>% unique
  setdiff(covid31_proteins, chromosome_export$`UniProt Entry`)
  covid31_AC <-
    chromosome_export %>% filter(`UniProt Entry` %in% covid31_proteins) %>%
select(`neXtProt AC`) %>% unlist(use.names = FALSE)

  rubtsova_proteins <-
    read_lines('data/raw_files/protein_lists/rubtsova.txt') %>% sort %>%
unique
  setdiff(rubtsova_proteins, chromosome_export$`UniProt Entry`)
  rubtsova_AC <-
    chromosome_export %>% filter(`UniProt Entry` %in% rubtsova_proteins) %>%
select(`neXtProt AC`) %>% unlist(use.names = FALSE)

  borchers_proteins <-
    read_lines('data/raw_files/protein_lists/borchers.txt') %>% sort %>%
unique
  setdiff(borchers_proteins, chromosome_export$`UniProt Entry`)
  borchers_AC <-
    chromosome_export %>% filter(`UniProt Entry` %in% borchers_proteins) %>%
select(`neXtProt AC`) %>% unlist(use.names = FALSE)

  top100_bpa <- read_tsv('data/raw_files/bpa_concentrations.txt')
  top100_bpa %<>% arrange(desc(`Concentration in fM/uL`)) %>% slice_head(n =
100L) %$% Entry
  setdiff(top100_bpa, chromosome_export$`UniProt Entry`)
  top100_AC <-
    chromosome_export %>% filter(`UniProt Entry` %in% top100_bpa) %>%
select(`neXtProt AC`) %>% unlist(use.names = FALSE)

  plasma_peptides <-
    as.character(
      Biostrings::readAAStringSet(
        'data/raw_files/peptideatlas/plasma/202304/APD_Hs_all.fasta'
      )
    )
```

```
  human_peptides <-
    as.character(
      Biostrings::readAAStringSet(
        'data/raw_files/peptideatlas/human/202401/APD_Hs_all.fasta'
      )
    )
  }

all_acc <-
  list(
    top100_bpa = top100_AC,
    borchers267 = borchers_AC,
    hbpp255 = hbpp255_AC,
    enc = enc_AC,
    covid31 = covid31_AC,
    rubtsova = rubtsova_AC
  )

priority <-
  tibble(
    list = c(
      'top100_bpa',
      'borchers267',
      'hbpp255',
      'enc',
      'covid31',
      'rubtsova'
    ),
    priority = c(10, 0, 0, 1000, 100, 100)
  )

protein_priority <- lapply(names(all_acc), function(n) {
  tibble(`neXtProt AC` = all_acc[[n]], list = n)
}) %>% bind_rows() %>% left_join(priority, by = 'list') %>% group_by(`neXtProt
AC`) %>% summarise(Priority = sum(priority))  %>% arrange(desc(Priority))

peptide_md5 <-
  peptide_sources %>% select(`Peptide Sequence`, md5) %>% unnest(md5) %>%
distinct
peptide_AC <-
  peptide_sources %>% select(`Peptide Sequence`, `neXtProt AC`) %>%
unnest(`neXtProt AC`) %>% distinct
AC_md5 <-
  isoform_sequences %>% select(`neXtProt AC`, md5) %>% distinct

peptide_AC <- as.data.table(peptide_AC)
```

```
peptide_md5 <- as.data.table(peptide_md5)
AC_md5 <- as.data.table(AC_md5)


gene_specific <-
  peptide_AC[, .(`Peptide Specific for Gene` = length(unique(`neXtProt AC`))
== 1L), `Peptide Sequence`]

gene_universal <-
  merge(
    merge(
      AC_md5,
      peptide_AC,
      by = 'neXtProt AC',
      all = TRUE,
      allow.cartesian = TRUE
    ),
    peptide_md5[, `Peptide in Isoform` := TRUE],
    by = c('Peptide Sequence', 'md5'),
    all = TRUE
  )[is.na(`Peptide in Isoform`), `Peptide in Isoform` := FALSE][, .(`Peptide
Present In Every Isoform` = all(`Peptide in Isoform`)), .(`neXtProt AC`,
`Peptide Sequence`)]

pepmap <-
  merge(gene_universal, gene_specific, by = 'Peptide Sequence')[, `Peptide
Length` := nchar(`Peptide Sequence`)]

pep_ct <-
  pepmap[, .(
    `Total Peptides` = uniqueN(.SD[`Peptide Length` >= 6L &
                                  `Peptide Length` <= 30L, `Peptide
Sequence`]),
    `Gene-specific Isoform-universal Peptides` = uniqueN(.SD[`Peptide Length`
>= 6L &
                                                               `Peptide
Length` <= 30L &
                                                               `Peptide
Present In Every Isoform` == TRUE &
                                                               `Peptide
Specific for Gene` == TRUE, `Peptide Sequence`])
  ), `neXtProt AC`]

pep_ct_per_gene_stat <-
  pep_ct[, .N, .(
    `Gene-specific Isoform-universal Peptides` = cut(
```

```
      `Gene-specific Isoform-universal Peptides`,
      breaks = c(-1L, 0L, 5L, 10L, 20L, 50L, 100L, 200L, 2500L),
      labels = c('0',
                 '1-5',
                 '6-10',
                 '11-20',
                 '21-50',
                 '51-100',
                 '101-200',
                 '> 200'),
      ordered = TRUE
    )
  )][order(`Gene-specific Isoform-universal Peptides`)]

fwrite(pep_ct_per_gene_stat, sep = '\t')

ggplot(pep_ct_per_gene_stat,
       aes(`Gene-specific Isoform-universal Peptides`, N)) +
geom_histogram(stat = 'identity') + theme_minimal(base_size = 18L) +
ggtitle('Gene-specific isoform-universal peptides of length 6-30 per gene')

cleaved_dt <- as.data.table(cleaved)

mod_ct <-
  cleaved_dt[, .(
    `Peptides 6-30` = .SD[`Peptide Length` >= 6L &
                            `Peptide Length` <= 30L, uniqueN(`Peptide
Sequence`)],
    `Phosphorylated Peptides` = .SD[`Peptide Length` >= 6L &
                                      `Peptide Length` <= 30L &
                                      `Phosphorylated Residues` > 0L,
uniqueN(`Peptide Sequence`)],
    `Modifed Peptides` = .SD[`Peptide Length` >= 6L &
                               `Peptide Length` <= 30L &
                               `Total Modified Residues` > 0L,
uniqueN(`Peptide Sequence`)]
  ), md5]
mod_ct[, `:=`(`Unmodified Peptides` = `Peptides 6-30` - `Modified Peptides`)]

mod_stat <- mod_ct[, .N, .(`Unmodified Peptides` = cut(
  `Unmodified Peptides`,
  breaks = c(-1L, 0L, 5L, 10L, 20L, 50L, 100L, 200L, 2500L),
  labels = c('0',
             '1-5',
             '6-10',
             '11-20',
```

```
              '21-50',
              '51-100',
              '101-200',
              '> 200'),
  ordered = TRUE
))][order(`Unmodified Peptides`)]

ggplot(mod_stat,
       aes(`Unmodified Peptides`, N)) + geom_histogram(stat = 'identity') +
theme_minimal(base_size = 18L) + ggtitle('Unmodified peptides of length 6-30
per isoform')

AC_md5[md5 %in% mod_ct[`Unmodified Peptides` == 0L, md5], uniqueN(`neXtProt
AC`)]
AC_md5[md5 %in% mod_ct[`Unmodified Peptides` > 0L & `Unmodified Peptides` <=
5L, md5], uniqueN(`neXtProt AC`)]

rag_ct <- cleaved_dt[, .(`Peptides with No Ragged Ends` = .SD[`Position
Penalty` < 1000L &
                                                                  `Peptide
Length` >= 6 &
                                                                  `Peptide
Length` <= 30, uniqueN(`Peptide Sequence`)]), md5]

rag_stat <- rag_ct[, .N, .(
                 `Peptides with No Ragged Ends` = cut(
                   `Peptides with No Ragged Ends`,
                   breaks = c(-1L, 0L, 5L, 10L, 20L, 50L, 100L, 200L, 2500L),
                   labels = c('0',
                              '1-5',
                              '6-10',
                              '11-20',
                              '21-50',
                              '51-100',
                              '101-200',
                              '> 200'),
                   ordered = TRUE
                 )
               )][order(`Peptides with No Ragged Ends`)]

ggplot(rag_stat,
       aes(`Peptides with No Ragged Ends`, N)) + geom_histogram(stat =
'identity') + theme_minimal(base_size = 18L) + ggtitle('Peptides with No
Ragged Ends of length 6-30 per isoform')

AC_md5[md5 %in% rag_ct[`Peptides with No Ragged Ends` == 0L, md5],
```

```
uniqueN(`neXtProt AC`)]
AC_md5[md5 %in% rag_ct[`Peptides with No Ragged Ends` > 0L & `Peptides with No
Ragged Ends` <= 5L, md5], uniqueN(`neXtProt AC`)]


peptides_dt <- as.data.table(peptides)

no_c <- peptides_dt[!`Has C` & `Peptide Length` >= 6 & `Peptide Length` <= 30,
unique(`Peptide Sequence`)]
pep_ct_c <- peptide_AC[, .(`Peptides without C` = .SD[`Peptide Sequence` %in%
no_c, .N]), `neXtProt AC`]

pep_c_stat <- pep_ct_c[, .N, .(
  `Peptides without C` = cut(
    `Peptides without C`,
    breaks = c(-1L, 0L, 5L, 10L, 20L, 50L, 100L, 200L, 2500L),
    labels = c('0',
               '1-5',
               '6-10',
               '11-20',
               '21-50',
               '51-100',
               '101-200',
               '> 200'),
    ordered = TRUE
  )
)][order(`Peptides without C`)]


clusterExport(cluster, c('peptide_AC', 'AC_md5', 'peptide_md5'))

peptide_mapping <-
  (protein_priority %>% arrange(desc(Priority)) %$% `neXtProt AC`)  %>%
pblapply(cl = cluster, function(acc) {
    pep_acc <- peptide_AC %>% filter(`neXtProt AC` == acc)
    prot_AC_md5 <- AC_md5 %>% filter(`neXtProt AC` == acc)
    pep_md5 <-
      peptide_md5 %>% filter(md5 %in% prot_AC_md5$md5) %>% mutate(`In md5` =
TRUE)

    shared <-
      peptide_AC %>% filter(`Peptide Sequence` %in% pep_acc$`Peptide
Sequence`,
                            `neXtProt AC` != acc) %>% select(`Peptide
Sequence`) %>% distinct %>% mutate(`Shared` = TRUE)
```

```
    universal <-
      cross_join(prot_AC_md5, pep_acc %>% select(`Peptide Sequence`)) %>%
left_join(pep_md5, by = c('md5', 'Peptide Sequence')) %>% mutate(`In md5` =
replace(`In md5`, is.na(`In md5`), FALSE)) %>% group_by(`Peptide Sequence`)
%>% summarize(Universal = all(`In md5`))

    result <-
      left_join(universal, shared, by = 'Peptide Sequence') %>%
mutate(`neXtProt AC` = acc) %>% mutate(Shared = replace(Shared, is.na(Shared),
FALSE))
    result
  }) %>% bind_rows

peptide_mapping <-
  accessions  %>% pblapply(cl = cluster, function(acc) {
    pep_acc <- peptide_AC %>% filter(`neXtProt AC` == acc)
    prot_AC_md5 <- AC_md5 %>% filter(`neXtProt AC` == acc)
    pep_md5 <-
      peptide_md5 %>% filter(md5 %in% prot_AC_md5$md5) %>% mutate(`In md5` =
TRUE)

    shared <-
      peptide_AC %>% filter(`Peptide Sequence` %in% pep_acc$`Peptide
Sequence`,
                            `neXtProt AC` != acc) %>% select(`Peptide
Sequence`) %>% distinct %>% mutate(`Shared` = TRUE)

    universal <-
      cross_join(prot_AC_md5, pep_acc %>% select(`Peptide Sequence`)) %>%
left_join(pep_md5, by = c('md5', 'Peptide Sequence')) %>% mutate(`In md5` =
replace(`In md5`, is.na(`In md5`), FALSE)) %>% group_by(`Peptide Sequence`)
%>% summarize(Universal = all(`In md5`))

    result <-
      left_join(universal, shared, by = 'Peptide Sequence') %>%
mutate(`neXtProt AC` = acc) %>% mutate(Shared = replace(Shared, is.na(Shared),
FALSE))
    result
  }) %>% bind_rows

peptide_mapping %<>% mutate(`Uniqueness Penalty` = Shared * 1000 +
(!Universal) * 1000)

position_penalty <-
  isoform_sequences %>% select(md5, `neXtProt AC`) %>% left_join(cleaved, by =
'md5') %>% group_by(`neXtProt AC`, `Peptide Sequence`) %>%
```

```
  summarise(
    `N-terminal Peptide` = any(`N-terminal Peptide`),
    `C-terminal Peptide` = any(`C-terminal Peptide`),
    `Next Site` = min(`Next Site`),
    `Previous Site` = min(`Previous Site`),
    `Aggregated Position Penalty` = `N-terminal Peptide` * 1000L +
      `C-terminal Peptide` * 1000L +
      (`Next Site` < 5L) +
      (`Previous Site` < 5L) +
      (`Next Site` < 3L) * 1000L +
      (`Previous Site` < 3L) * 1000L,
    `Lowest Position Penalty` = min(`Position Penalty`),
    `Highest Position Penalty` = max(`Position Penalty`),
    `Modification Penalty` = max(`Modification Penalty`),
    `Variant Penalty` = max(`Variant Penalty`),
    `Times seen` = n()
  )

peptide_scores <-
  protein_priority %>% left_join(peptide_mapping, by = 'neXtProt AC') %>%
left_join(peptides, by = 'Peptide Sequence') %>% left_join(position_penalty,
by = c('neXtProt AC', 'Peptide Sequence')) %>% mutate(
    `Total Penalty` = `Uniqueness Penalty` + `Composition Penalty` +
`Aggregated Position Penalty` + `Modification Penalty` + `Variant Penalty`
  )

peptide_scores %>% select(
  `neXtProt AC`,
  Priority,
  `Peptide Sequence`,
  `Total Penalty`,
  `Uniqueness Penalty`,
  `Composition Penalty`,
  `Aggregated Position Penalty`,
  `Modification Penalty`,
  `Variant Penalty`
)

peptide_scores %<>% mutate(`Borchers267 Protein` = `neXtProt AC` %in%
borchers_AC)
peptide_scores %<>% mutate(`Borchers267 Peptide` = `Peptide Sequence` %in%
borchers_peptides)
peptide_scores %<>% mutate(`HBPP255 Peptide` = `Peptide Sequence` %in%
hbpp255_peptides)
peptide_scores %<>% mutate(`Peptide Atlas Human Plasma` = `Peptide Sequence`
%in% plasma_peptides)
```

```
peptide_scores %<>% mutate(`Peptide Atlas Human All` = `Peptide Sequence` %in%
human_peptides)

peptides %<>% mutate(`Peptide Atlas Human Plasma` = `Peptide Sequence` %in%
plasma_peptides)
peptides %<>% mutate(`Peptide Atlas Human All` = `Peptide Sequence` %in%
human_peptides)

ggplot(peptide_scores %>% filter(`Borchers267 Protein`),
       aes(`Total Penalty`)) + geom_histogram(binwidth = 1L, center = 0L) +
facet_grid(`Borchers267 Peptide` ~

., scales = 'free_y')

peptide_scores %>% filter(`Total Penalty` == 0) %$% `neXtProt AC` %>% unique
%>% length

selected_peptides_1 <- peptide_scores %>%
  filter(
    Priority >= 1000,
    `Peptide Length` >= 8L,
    `Peptide Length` <= 20L,
    `Borchers267 Peptide` |
      `Peptide Atlas Human Plasma` | `Peptide Atlas Human All`,
    `Total Penalty` < 1000L
  ) %>%
  group_by(`neXtProt AC`) %>%
  slice_min(`Total Penalty`, with_ties = FALSE) %>%
  select(
    `neXtProt AC`,
    `Peptide Sequence`,
    `Peptide Length`,
    `Total Penalty`,
    `Borchers267 Peptide`,
    `Peptide Atlas Human Plasma`,
    `Peptide Atlas Human All`
  ) %>%
  mutate(`Peptide Set` = 1L)

selected_peptides_2 <- peptide_scores %>%
  filter(
    Priority >= 1000,
    `Peptide Length` >= 8L,
    `Peptide Length` <= 20L,
    !(`neXtProt AC` %in% selected_peptides_1$`neXtProt AC`),
    `Total Penalty` < 1000L
```

```
  ) %>%
  group_by(`neXtProt AC`) %>%
  slice_min(`Total Penalty`, with_ties = FALSE) %>%
  select(
    `neXtProt AC`,
    `Peptide Sequence`,
    `Peptide Length`,
    `Total Penalty`,
    `Borchers267 Peptide`,
    `Peptide Atlas Human Plasma`,
    `Peptide Atlas Human All`
  ) %>%
  mutate(`Peptide Set` = 2L)

selected_peptides_3 <- peptide_scores %>%
  filter(
    Priority >= 1000,
    `Peptide Length` >= 6L,
    `Peptide Length` <= 30L,
    !(`neXtProt AC` %in% selected_peptides_1$`neXtProt AC`),
    !(`neXtProt AC` %in% selected_peptides_2$`neXtProt AC`),
    `Total Penalty` < 1000L
  ) %>%
  group_by(`neXtProt AC`) %>%
  slice_min(`Total Penalty`, with_ties = FALSE) %>%
  select(
    `neXtProt AC`,
    `Peptide Sequence`,
    `Peptide Length`,
    `Total Penalty`,
    `Borchers267 Peptide`,
    `Peptide Atlas Human Plasma`,
    `Peptide Atlas Human All`
  ) %>%
  mutate(`Peptide Set` = 3L)

selected_peptides <-
  list(selected_peptides_1,
       selected_peptides_2,
       selected_peptides_3) %>% bind_rows()

fwrite(selected_peptides,
       'peptides_20240312.txt',
       sep = '\t',
       eol = '\n')
```

```
selected_peptides_4 <- peptide_scores %>%
  filter(
    Priority < 1000,
    Priority >= 100,
    `Peptide Length` >= 8L,
    `Peptide Length` <= 20L,
    `Borchers267 Peptide` |
      `Peptide Atlas Human Plasma` | `Peptide Atlas Human All`,
    !(`neXtProt AC` %in% selected_peptides$`neXtProt AC`),
    `Total Penalty` < 1000L
  ) %>%
  group_by(`neXtProt AC`) %>%
  slice_min(`Total Penalty`, with_ties = FALSE) %>%
  select(
    `neXtProt AC`,
    `Peptide Sequence`,
    `Peptide Length`,
    `Total Penalty`,
    `Borchers267 Peptide`,
    `Peptide Atlas Human Plasma`,
    `Peptide Atlas Human All`
  ) %>%
  mutate(`Peptide Set` = 1L)

selected_peptides_5 <- peptide_scores %>%
  filter(
    Priority >= 1000,
    Priority >= 100,
    `Peptide Length` >= 8L,
    `Peptide Length` <= 20L,
    !(`neXtProt AC` %in% selected_peptides$`neXtProt AC`),
    !(`neXtProt AC` %in% selected_peptides_4$`neXtProt AC`),
    `Total Penalty` < 1000L
  ) %>%
  group_by(`neXtProt AC`) %>%
  slice_min(`Total Penalty`, with_ties = FALSE) %>%
  select(
    `neXtProt AC`,
    `Peptide Sequence`,
    `Peptide Length`,
    `Total Penalty`,
    `Borchers267 Peptide`,
    `Peptide Atlas Human Plasma`,
    `Peptide Atlas Human All`
  ) %>%
```

```
  mutate(`Peptide Set` = 2L)

selected_peptides_6 <- peptide_scores %>%
  filter(
    Priority >= 1000,
    Priority >= 100,
    `Peptide Length` >= 6L,
    `Peptide Length` <= 30L,
    !(`neXtProt AC` %in% selected_peptides$`neXtProt AC`),
    !(`neXtProt AC` %in% selected_peptides_4$`neXtProt AC`),
    !(`neXtProt AC` %in% selected_peptides_5$`neXtProt AC`),
    `Total Penalty` < 1000L
  ) %>%
  group_by(`neXtProt AC`) %>%
  slice_min(`Total Penalty`, with_ties = FALSE) %>%
  select(
    `neXtProt AC`,
    `Peptide Sequence`,
    `Peptide Length`,
    `Total Penalty`,
    `Borchers267 Peptide`,
    `Peptide Atlas Human Plasma`,
    `Peptide Atlas Human All`
  ) %>%
  mutate(`Peptide Set` = 3L)

selected_peptides_v2 <-
  list(selected_peptides_4,
       selected_peptides_5,
       selected_peptides_6) %>% bind_rows()

fwrite(selected_peptides_v2,
       'peptides_20240312_v2.txt',
       sep = '\t',
       eol = '\n')
```

#### Script for peptide scoring

```
# Script to collect and store information about proteins from neXtProt via
REST API and tidyverse
# This script modifies neXtProt responses to allow caching them for a year!

library(magrittr)
library(httr2)
```

```
library(dplyr)
library(readr)
library(tidyr)
library(stringi)
library(Peptides)
library(pbapply)
library(parallel)
library(data.table)
library(ggplot2)

# library(dtplyr)

options(readr.show_col_types = FALSE)
nextprot <-
  readRDS('data/nextprot_rest_api_cache/2023-09-11_2.40.0/neXtProt.rds')

# Compute peptide penalties based on peptide's position in a chain, PTMs or
Sequence Variants

nextprot$cleaved %<>% mutate(
  `Position Penalty` =  `N-terminal Peptide` * 1000L +
    `C-terminal Peptide` * 1000L +
    (`Next Site` < 5L) +
    (`Previous Site` < 5L) +
    (`Next Site` < 3L) * 1000L +
    (`Previous Site` < 3L) * 1000L,
  `Modification Penalty` = `Total Modified Residues` * 1000L,
  `Variant Penalty` = `Sequence Variants` * 1000L
)

# Compute peptide penalty based on it's amino acid composition

nextprot$peptides %<>% mutate(
  `Composition Penalty` =
    `Has C` * 1000L + # Prohibited (standards should be modified)
    `Has W` * 1000L + # Prohibited (oxidizes heavily with several channels)
    `Has N` + # Deamidation
    `Has Q` + # Deamidation
    `Has M` + # Oxidation
    `Has ^E` * 1000L + # These N and Q are penalized additionaly, they are
prone to cyclization during standard synthesis
    `Has ^Q` * 1000L + # These N and Q are penalized additionaly, they are
prone to cyclization during standard synthesis
    `Has NG` + # These N and Q are penalized additionaly, they are more prone
to modification
    `Has NP` + # These N and Q are penalized additionaly, they are more prone
```

```
to modification
    `Has DP` + # Unstable
    `Has KP` + # This cleavage occurs but slowly
    `Has RP` + # This cleavage occurs but slowly
    `Has O` * 1000L + # Prohibited (synthesis / biosynthesis problems)
    `Has X` * 1000L + # Prohibited (synthesis / biosynthesis problems)
    `Has U` * 1000L + # Prohibited (synthesis / biosynthesis problems)
    `Longest Repeat` - 1L + # Consecutive identical residues are sometimes
difficult in synthesis
    as.integer(`Peptide Length` < 6L) * 1000L + # Short are less specific and
too hydrophylic
    as.integer(`Peptide Length` < 8L) + # Short are less specific
    as.integer(`Peptide Length` > 30) * 1000L + # Long are less stable and
more expensive
    as.integer(is.na(`m/z +2`) || is.na(`m/z +3`) || (`m/z +2` < 300 &
       `m/z +3` < 300)) * 1000L + # Below 300 Da the background is high and MS
less stable
    as.integer(is.na(`m/z +2`) || is.na(`m/z +3`) || (`m/z +2` > 1500 &
       `m/z +3` > 1500)) * 1000L + # Above 1500 Da MS less stable
    as.integer(is.na(`Hydrophobicity`) || (Hydrophobicity < -2) +
is.na(`Hydrophobicity`) || (Hydrophobicity > 1)) # Early and late peptides
depend too much on stationary phase properties
)

nextprot$peptide_mapping %<>% mutate(
  `Uniqueness Penalty` = (!`Peptide Specific for Gene`) * 1000 + (!`Peptide
Present In Every Isoform`) * 1000
)

nextprot$position_penalty <- nextprot$isoform_sequences %>% select(md5,
`neXtProt AC`) %>%
  left_join(nextprot$cleaved, by = 'md5', relationship = 'many-to-many') %>%
  group_by(`neXtProt AC`, `Peptide Sequence`) %>%
  summarise(
    `N-terminal Peptide` = any(`N-terminal Peptide`),
    `C-terminal Peptide` = any(`C-terminal Peptide`),
    `Next Site` = min(`Next Site`),
    `Previous Site` = min(`Previous Site`),
    `Aggregated Position Penalty` = `N-terminal Peptide` * 1000L +
      `C-terminal Peptide` * 1000L +
      (`Next Site` < 5L) +
      (`Previous Site` < 5L) +
      (`Next Site` < 3L) * 1000L +
      (`Previous Site` < 3L) * 1000L,
    `Lowest Position Penalty` = min(`Position Penalty`),
    `Highest Position Penalty` = max(`Position Penalty`),
```

```
    `Modification Penalty` = max(`Modification Penalty`),
    `Variant Penalty` = max(`Variant Penalty`)
  )

peptide_mapping <- as.data.table(nextprot$peptide_mapping)[, .(
  `neXtProt AC` = paste0(`neXtProt AC`, collapse = '; '),
  `Protein Count` = length(unique(`neXtProt AC`)),
  `Peptide Present In Every Isoform` = all(`Peptide Present In Every
Isoform`),
  `Peptide Specific for Gene` = all(`Peptide Specific for Gene`),
  `Uniqueness Penalty` = max(`Uniqueness Penalty`)
), .(`Peptide Sequence`)]

peptide_composition <- as.data.table(nextprot$peptides)

peptide_position <- as.data.table(nextprot$position_penalty)[, .(
  `Max Aggregated Position Penalty` = max(`Aggregated Position Penalty`),
  `Max Variant Penalty` = max(`Variant Penalty`),
  `Max Modification Penalty` = max(`Modification Penalty`),
  `N-terminal Peptide` = any(`N-terminal Peptide`),
  `C-terminal Peptide` = any(`C-terminal Peptide`),
  `Next Site` = min(`Next Site`),
  `Previous Site` = min(`Previous Site`)
), .(`Peptide Sequence`)]

peptide_scores <- merge(
  merge(peptide_mapping, peptide_composition, by = 'Peptide Sequence'),
  peptide_position,
  by = 'Peptide Sequence'
)

fwrite(peptide_scores, 'peptide_scores.txt')
```
